# Coherent feedforward regulation of gene expression by *Caulobacter* σ^T^ and GsrN during hyperosmotic stress

**DOI:** 10.1101/344820

**Authors:** Matthew Z. Tien, Benjamin J. Stein, Sean Crosson

## Abstract

GsrN is a conserved small RNA that is under transcriptional control of the general stress sigma factor, σ^T^, and that functions as a post-transcriptional regulator of *Caulobacter crescentus* survival under multiple stress conditions. We have defined features of GsrN structure that determine survival under hyperosmotic stress, and have applied transcriptomic and proteomic methods to identify regulatory targets of GsrN under hyperosmotic conditions. The 5’ end of GsrN, which includes a conserved cytosine-rich stem loop structure, is necessary for cell survival after osmotic upshock. GsrN both activates and represses gene expression in this stress condition. Expression of an uncharacterized open reading frame predicted to encode a glycine-zipper protein, *osrP*, is strongly activated by GsrN. Our data support a model in which GsrN physically interacts with *osrP* mRNA through its 5’ C-rich stem loop to enhance OsrP protein expression. We conclude that *sigT*, *gsrN*, and *osrP* form a coherent feedforward loop in which σ^T^ activates *gsrN* and *osrP* transcription during stress, and GsrN activates OsrP protein expression at the post-transcriptional level. This study delineates transcriptional and post-transcriptional layers of *Caulobacter* gene expression control during hyperosmotic stress, uncovers a new regulatory target of GsrN, and defines a coherent feedforward motif in the *Caulobacter* GSR regulatory network.

**Importance:** Bacteria inhabit diverse niches, and must adapt their physiology to constant environmental fluctuations. A major response to environmental perturbation is to change gene expression. *Caulobacter* and other alphaproteobacteria initiate a complex gene expression program known as the general stress response (GSR) under conditions including oxidative stress, osmotic stress, and nutrient limitation. The GSR enables cell survival in these environments. Understanding how bacteria survive stress requires that we dissect gene expression responses, such as the GSR, at the molecular level. This study is significant as it defines transcriptional and post-transcriptional layers of gene expression regulation in response to hyperosmotic stress. We further provide evidence that coherent feedforward motifs influence the system properties of the *Caulobacter* GSR pathway.

## Introduction

Cells alter gene expression to adapt to environmental perturbations. In bacteria, two major mechanisms controlling transcription are two-component signaling (TCS) (1) and alternative sigma (σ) factor regulation (2, 3). In species of the class Alphaproteobacteria, crosstalk between these mechanisms is uniquely achieved via the protein, PhyR, which contains both a σ-like domain and a TCS receiver domain (4–7). Under a range of specific stress conditions, PhyR becomes phosphorylated and, through a protein partner switching mechanism (5), activates a gene expression program known as the general stress response (GSR). The GSR is required for survival under diverse environmental conditions (8, 9).

We recently developed a network-based algorithm (10) to interrogate publicly available gene expression datasets to predict genes functioning in stress survival in the alphaproteobacterium, *Caulobacter crescentus*. This led to the discovery of a conserved small RNA (sRNA), GsrN, that plays an important role in survival across distinct environmental conditions including hyperosmotic and oxidative stress (11). GsrN is directly activated by the GSR alternative sigma factor, σ^T^, and imposes a post-transcriptional layer of gene expression regulation during the general stress response. In the case of hydrogen peroxide stress, GsrN protects cells by base pairing with the 5’ leader sequence of *katG* mRNA to promote expression of KatG, a catalase/peroxidase protein (11). To date, the identity of genes regulated by GsrN under hyperosmotic stress conditions remain undefined. The goal of this study was to define structural features of *Caulobacter* GsrN that are required for hyperosmotic stress survival and to identify direct molecular targets of GsrN under hyperosmotic conditions.

The induction of sRNA expression by osmotic stress has been described in a handful of bacterial species (12–15). Examples of sRNAs with known roles in osmoregulation of gene expression include OmrA/OmrB (16), MicF (12), and MicC (17) in *Escherichia coli*. The OmrA/OmrB system is upregulated during osmotic stress by the two-component system, EnvZ-OmpR. OmrA/OmrB function as post-transcriptional feedback repressors of OmpR (18) and repress the expression of outer membrane proteins, including TonB-dependent receptors (16). MicF and MicC are also induced by changes in osmolarity and function to repress translation of outer membrane proteins OmpF and OmpC, respectively (12, 17). Though expression of these sRNAs are induced by shifts in the osmotic state of the environment, data demonstrating roles for OmrA/OmrB, MicC, and MicF in cell survival under acute osmotic stress have not, to our knowledge, been reported.

We have assayed hyperosmotic stress survival of a series of *Caulobacter gsrN* mutant strains, used transcriptomic and proteomic methods to more clearly define the role of GsrN in gene expression during hyperosmotic stress, and provided evidence for a new direct regulatory target of GsrN. Features of GsrN structure that are functionally important for hyperosmotic stress survival are contained in the 5’ end of the molecule, and include a conserved cytosine-rich stem loop structure. Transcriptomic and proteomic analyses identified genes that are both activated and repressed by GsrN upon shift to a hyperosmotic environment. Among the regulated gene set was a hypothetical open reading frame we have named *osrP*, which encodes a glycine zipper domain resembling the glycine zipper motifs of large- and small-conductance mechanosensitive channels (19). We present evidence that GsrN directly interacts with *osrP* mRNA and activates OsrP protein expression at the post-transcriptional level to form a coherent feedforward regulatory loop with σ^T^. This study advances understanding of *Caulobacter crescentus* gene expression during hyperosmotic stress and defines a new post-transcriptional regulatory target of GsrN.

## Results

### A 5’ cytosine-rich loop in GsrN is necessary for osmotic stress survival

GsrN is a small RNA (sRNA) that undergoes endonucleolytic processing, and functions as a potent regulator of both oxidative stress and osmotic stress survival in *Caulobacter crescentus* (11). Expression of the processed 5’ fragment of GsrN is necessary and sufficient to protect cells from hydrogen peroxide exposure. This protection requires interaction of GsrN with the mRNA of catalase/peroxidase *katG* through a C-rich loop located in the stable 5’ half of GsrN (11). To assess the functional role of GsrN processing and the 5’ C-rich loop under a distinct stress condition, we assayed osmotic stress survival of strains harboring truncated and C-loop mutant variants of GsrN.

For these assays, we generated: *i)* a GsrN deletion strain (Δ*gsrN*), and *ii)* a strain lacking the 5’ end of GsrN, *gsrN*(Δ5’) (by deleting *gsrN* nucleotides 10–50) (**Fig. 1A**). Both Δ*gsrN* and *gsrN*(Δ5’) had ≈1 order of magnitude reduced viability during sucrose-induced osmotic stress when compared to wild-type *Caulobacter* strain CB15 **(Fig. 1B**). Ectopic expression of the first 58 nucleotides of *gsrN* in single copy from its native promoter (*gsrN*(Δ3’)) complemented the survival defect of Δ*gsrN*. Notably, a Δ*gsrN* strain harboring multiple integrations of this complementation plasmid, Δ*gsrN*::*gsrN*(Δ3’)^++^, had increased viability under hyperosmotic stress compared to wild type (**Fig. 1B**). This protective effect is consistent with peroxide stress protection conferred by full-length *gsrN* overexpression (*gsrN*^++^), reported in our previous study (11).

**FIG 1.**
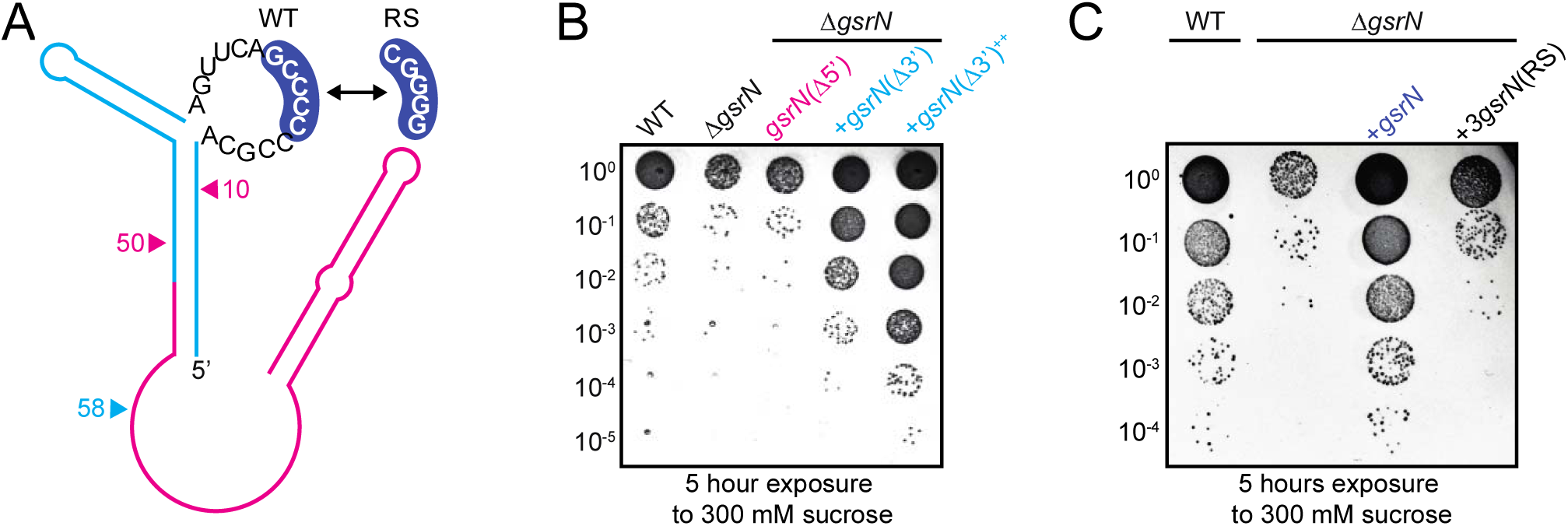
Modifying the 5’ cytosine-rich loop of GsrN reduces *Caulobacter* viability under hyperosmotic stress. (A) Secondary structure model of GsrN. Bases in the 5’ C-rich loop are displayed. GsrN undergoes endonucleolytic processing; cyan lines indicate the 5’ end of GsrN and pink lines indicate the 3’ end of GsrN (post-processing). Pink arrows refer to residues 10 and 50, which are the sites of deletion in the strain *gsrN*(Δ5’). Cyan arrow marks the 5’ end of GsrN construct, *gsrN*(Δ3’). Blue highlighted bases in the C-rich loop of GsrN were replaced in the mutant, *gsrN*(RS). (B) Hyperosmotic stress survival assay of *Caulobacter* wild type (WT) and *gsrN* mutant strains. Strains were treated with 300 mM sucrose for 5 hours and colony forming units (CFUs) were enumerated. (C) Hyperosmotic survival assay of WT and Δ*gsrN* complemented with either wild-type *gsrN* or *gsrN(RS).* Plates in B and C are representative of triplicate assays. Quantification of CFUs in treated versus untreated strains are presented in Fig. S1 in supplemental material.

Considering expression of the 5’ end of GsrN complemented the hyperosmotic stress survival defect of Δ*gsrN*, we hypothesized that the 5’ C-rich loop functions to mitigate osmotic stress in addition to its previously reported function in peroxide stress mitigation. Overexpression of a GsrN mutant variant in which the 5’ cytosine tract was replaced with guanosines, *gsrN*(RS), failed to restore osmotic stress survival to wild-type levels in the Δ*gsrN* strain (**Fig. 1C**). We thus propose that the 5’ C-loop of GsrN is necessary to target mRNAs that are involved in osmotic stress survival.

### *gsrN*-dependent osmotic stress protection requires *sigT*

*gsrN* expression is directly activated by the general stress sigma factor, SigT (σ^T^). As described above, strains lacking *sigT* (6, 20) or *gsrN* (11) are more susceptible to hyperosmotic stress. We thus tested whether expression of *gsrN* is sufficient to rescue the osmotic stress survival defect of the Δ*sigT* strain, as previously reported for hydrogen peroxide stress (11). We constructed a Δ*sigT* strain in which *gsrN* transcription was driven by the primary sigma factor RpoD (σ^70^). We call this expression system P1-*gsrN* (**Fig. 2A**). Expression of GsrN from P1 resulted in comparable steady-state levels to GsrN expressed from its native σ^T^-dependent promoter (**Fig. 2B**), but did not rescue the hyperosmotic stress survival defect of Δ*sigT* (**Fig. 2C**). Unlike acute peroxide stress, which does not induce expression of *gsrN*, osmotic stress induces the *gsrN* transcription by a factor of three (11). To better emulate GsrN expression during osmotic stress, we created a strain bearing three copies of P1-*gsrN*. Using this 3(P1-*gsrN*) strain, we matched the enhanced steady-state levels of GsrN observed during osmotic stress (**Fig. 2B**). However, enhanced expression of GsrN in Δ*sigT*+3(P1-*gsrN*) still failed to rescue the hyperosmotic stress survival defect of the Δ*sigT* strain (**Fig. 2C**). We conclude that GsrN-dependent protection during hyperosmotic stress requires other genes in the σ^T^-regulon.

**FIG 2.**
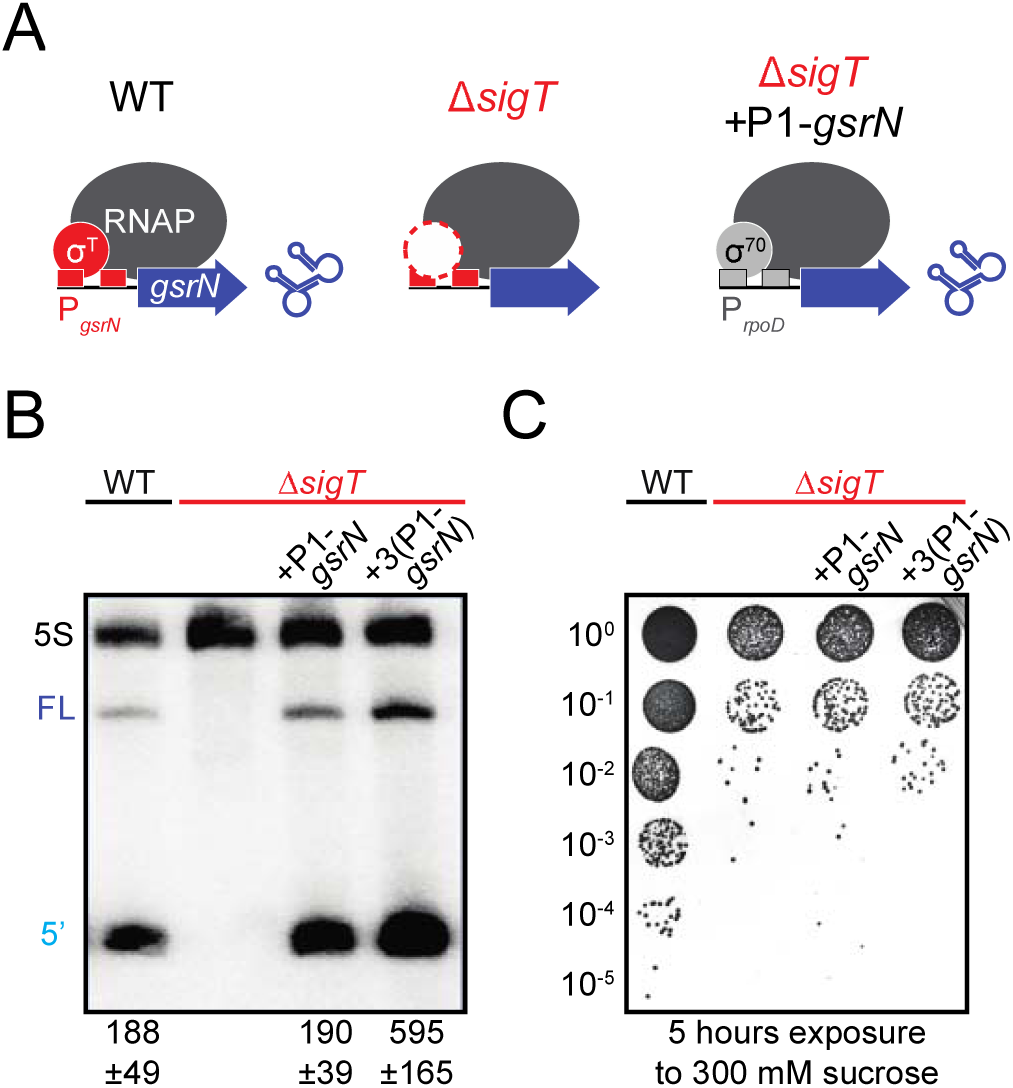
*gsrN*-dependent osmotic stress protection requires *sigT*. (A) Schematic of *gsrN* transcription in wild type (WT), Δ*sigT*, and a Δ*sigT* strain bearing the P1-*gsrN* expression system (Δ*sigT*+P1-*gsrN*). P1 is a RpoD(σ^70^)-dependent promoter. (B) Northern blot of total RNA from wild type, Δ*sigT*, Δ*sigT*+P1-*gsrN*, and Δ*sigT*+3(P1-*gsrN*) probed with radiolabeled oligos specific to GsrN and 5S rRNA (loading control). Labels on the left refer to 5S rRNA (5S in black), full-length GsrN (FL in dark blue), and the 5’ isoform of GsrN (5’ in cyan). Quantified values below the blot are mean ± SD of total (FL + 5’) normalized signal, n = 3 independent replicates. (C) Hyperosmotic stress survival assay of the strains in B. Plate is representative of triplicate assays. Quantification of CFUs in treated versus untreated strains are presented in Fig. S1 in supplemental material.

### Defining *sigT* and *gsrN* regulated genes under hyperosmotic conditions

We considered two non-mutually exclusive models to explain why GsrN-dependent protection against hyperosmotic stress requires σ^T^: *i)* GsrN functions as a direct post-transcriptional regulator of mRNAs that are transcribed by σ^T^, *ii)* GsrN regulates gene products that are not under the control of σ^T^, but that require σ^T^-regulated genes to mitigate hyperosmotic stress. Thus, to identify candidate mRNA targets and begin gathering evidence to support either (or both) models, we measured gene expression during hyperosmotic stress in a GsrN overexpression strain (*gsrN*^++^), a Δ*sigT* strain, and in wild type *Caulobacter*. Specifically, we measured steady-state transcript levels in Δ*sigT* and wild type strains under stressed and untreated conditions to define the σ^T^-dependent osmotic stress regulon. We further measured transcripts in *gsrN*^++^ and wild type under the same conditions to identify candidate transcripts involved in *gsrN*-dependent hyperosmotic stress protection. Lastly, we measured proteome changes between treated and untreated *gsrN*^++^ and wild-type strains to define protein expression regulated by GsrN during osmotic stress.

Although *sigT*-dependent gene expression has been previously studied in *Caulobacter* using microarray technologies (7, 20, 21), a high-resolution RNA-seq analysis of GSR mutant strains under hyperosmotic stress has not been published. Our RNA-seq measurements defined a σ^T^-regulon comprising 333 genes that are differentially expressed between Δ*sigT* and wild type under untreated conditions (false-discovery rate (FDR) *p-value* ≤0.05; absolute fold change ≥ 1.5). The number of differentially regulated genes during hyperosmotic stress is greater – 530 genes – using the same cutoff criteria. We defined the core σ^T^-regulon as the intersection of differentially regulated genes in both untreated and treated conditions, 220 genes **(Table S3)**. This expands the number of σ^T^-regulated genes compared to previous reports by our group (20) and others (21). We further sought to predict genes in the σ^T^ regulon that are directly transcribed by σ^T^. To this end, we extracted 250 nucleotide windows upstream of the translation start sites of genes activated by *sigT*. In the case of operons, we only considered the upstream region of the leading gene. We then created a degenerate σ^T^ motif based on variations in 20 previously identified σ^T^-binding sites **(Fig. S2)** (7, 20, 21). We searched for this motif in the regions upstream of *sigT*-activated genes and identified 32 additional transcripts with candidate σ^T^-binding sites **(see Table S3)**.

A parallel RNA-Seq experiment identified 35 genes that are differentially expressed in *gsrN*^++^ relative to wild type in untreated conditions and 141 genes under hyperosmotic conditions (false-discovery rate (FDR) *p-value* ≤0.05; absolute fold change ≥ 2.0) **(see Table S4)**. Considering that differences in GsrN-regulated transcripts do not necessarily correspond to differences in protein levels (11), and that GsrN is known to regulate gene expression at the post-transcriptional level, we performed a LC-MS/MS analysis of total soluble protein isolated from *gsrN*^++^ and wild type strains under untreated and hyperosmotic conditions. Twenty-two proteins showed significant differences in steady-state levels between *gsrN*^++^ and wild type under untreated conditions (false-discovery rate (FDR) *p-value* ≤0.05; absolute fold change ≥ 2.0). None of these proteins showed significant transcript level differences under the same criteria, and in all cases protein levels were lower in *gsrN*^++^ strains compared to wild type in untreated conditions **(Table 1)**. This provides evidence that the predominant role for *gsrN* in exponentially-growing cells is a repressor. Under hyperosmotic stress nine proteins had significant differences in steady-state levels between *gsrN*^++^ and wild type **(Table 2)**. Four of these proteins had corresponding significant differences in transcript levels; one protein had an inverse relationship with its transcript levels (**Fig. 3**). This analysis identified proteins for which expression is activated by GsrN under hyperosmotic stress and proteins for which expression is repressed by GsrN under hyperosmotic stress.

**TABLE 1.**
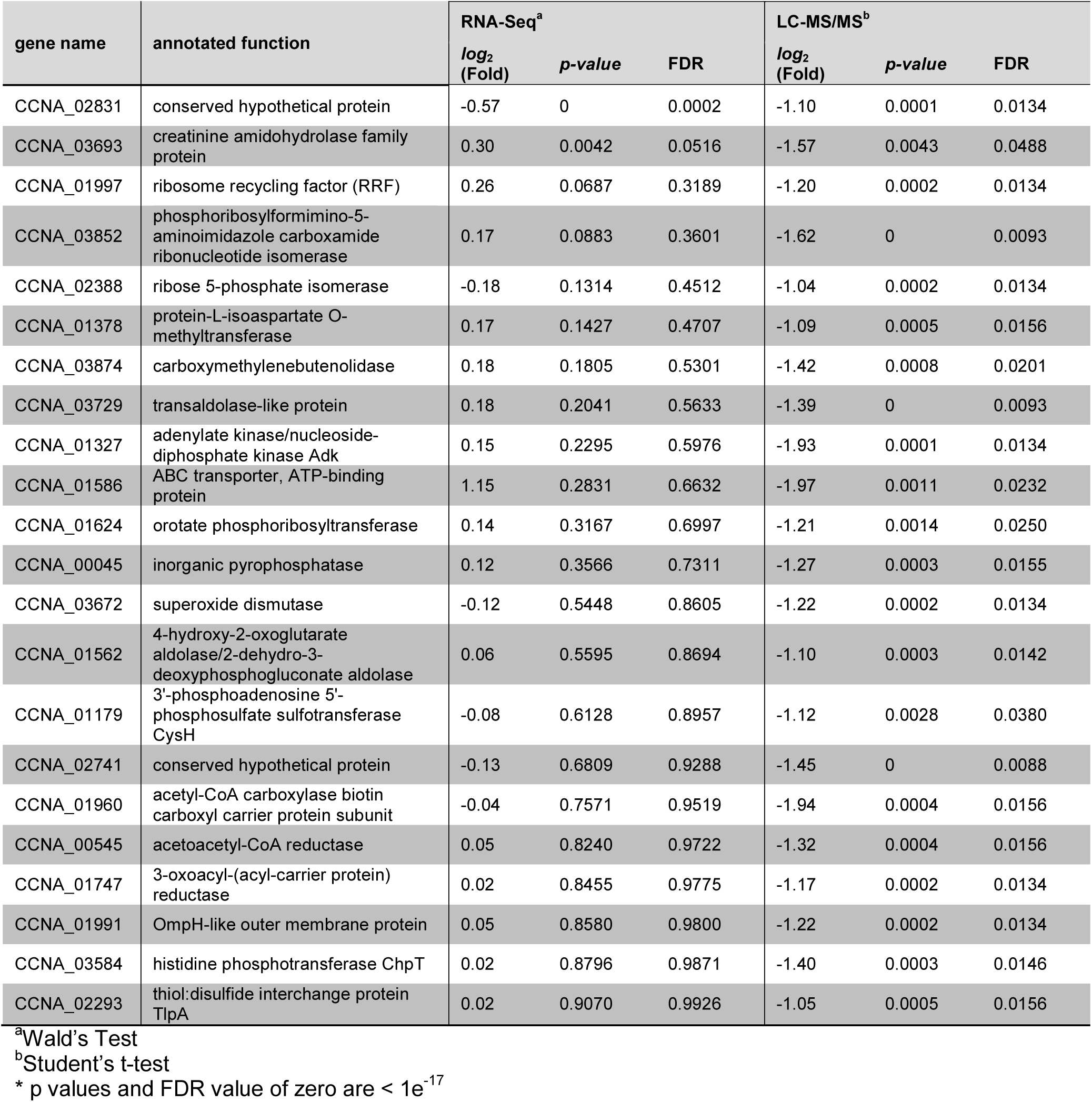
Proteins with significant differences in steady-state levels between *gsrN*^++^ and wild type (*gsrN*^++^ / WT), with associated transcript changes.

| gene name | annotated function | RNA-Seq<sup>a</sup> |  |  | LC-MS/MS<sup>b</sup> |  |  |
| --- | --- | --- | --- | --- | --- | --- | --- |
|  |  | *log<sub>2</sub>* (Fold) | *p*-value | FDR | *log<sub>2</sub>* (Fold) | *p*-value | FDR |
| CCNA_02831 | conserved hypothetical protein | -0.57 | 0 | 0.0002 | -1.10 | 0.0001 | 0.0134 |
| CCNA_03693 | creatinine amidohydrolase family protein | 0.30 | 0.0042 | 0.0516 | -1.57 | 0.0043 | 0.0488 |
| CCNA_01997 | ribosome recycling factor (RRF) | 0.26 | 0.0687 | 0.3189 | -1.20 | 0.0002 | 0.0134 |
| CCNA_03852 | phosphoribosylformimino-5-aminoimidazole carboxamide ribonucleotide isomerase | 0.17 | 0.0883 | 0.3601 | -1.62 | 0 | 0.0093 |
| CCNA_02388 | ribose 5-phosphate isomerase | -0.18 | 0.1314 | 0.4512 | -1.04 | 0.0002 | 0.0134 |
| CCNA_01378 | protein-L-isoaspartate O-methyltransferase | 0.17 | 0.1427 | 0.4707 | -1.09 | 0.0005 | 0.0156 |
| CCNA_03874 | carboxymethylenebutenolidase | 0.18 | 0.1805 | 0.5301 | -1.42 | 0.0008 | 0.0201 |
| CCNA_03729 | transaldolase-like protein | 0.18 | 0.2041 | 0.5633 | -1.39 | 0 | 0.0093 |
| CCNA_01327 | adenylate kinase/nucleoside-diphosphate kinase Adk | 0.15 | 0.2295 | 0.5976 | -1.93 | 0.0001 | 0.0134 |
| CCNA_01586 | ABC transporter, ATP-binding protein | 1.15 | 0.2831 | 0.6632 | -1.97 | 0.0011 | 0.0232 |
| CCNA_01624 | orotate phosphoribosyltransferase | 0.14 | 0.3167 | 0.6997 | -1.21 | 0.0014 | 0.0250 |
| CCNA_00045 | inorganic pyrophosphatase | 0.12 | 0.3566 | 0.7311 | -1.27 | 0.0003 | 0.0155 |
| CCNA_03672 | superoxide dismutase | -0.12 | 0.5448 | 0.8605 | -1.22 | 0.0002 | 0.0134 |
| CCNA_01562 | 4-hydroxy-2-oxoglutarate aldolase/2-dehydro-3-deoxyphosphogluconate aldolase | 0.06 | 0.5595 | 0.8694 | -1.10 | 0.0003 | 0.0142 |
| CCNA_01179 | 3'-phosphoadenosine 5'-phosphosulfate sulfotransferase CysH | -0.08 | 0.6128 | 0.8957 | -1.12 | 0.0028 | 0.0380 |
| CCNA_02741 | conserved hypothetical protein | -0.13 | 0.6809 | 0.9288 | -1.45 | 0 | 0.0088 |
| CCNA_01960 | acetyl-CoA carboxylase biotin carboxyl carrier protein subunit | -0.04 | 0.7571 | 0.9519 | -1.94 | 0.0004 | 0.0156 |
| CCNA_00545 | acetoacetyl-CoA reductase | 0.05 | 0.8240 | 0.9722 | -1.32 | 0.0004 | 0.0156 |
| CCNA_01747 | 3-oxoacyl-(acyl-carrier protein) reductase | 0.02 | 0.8455 | 0.9775 | -1.17 | 0.0002 | 0.0134 |
| CCNA_01991 | OmpH-like outer membrane protein | 0.05 | 0.8580 | 0.9800 | -1.22 | 0.0002 | 0.0134 |
| CCNA_03584 | histidine phosphotransferase ChpT | 0.02 | 0.8796 | 0.9871 | -1.40 | 0.0003 | 0.0146 |
| CCNA_02293 | thiol:disulfide interchange protein TlpA | 0.02 | 0.9070 | 0.9926 | -1.05 | 0.0005 | 0.0156 |

<sup>a</sup>Wald's Test <sup>b</sup>Student's t-test \* *p* values and FDR value of zero are < 1e<sup>-17</sup>

**TABLE 2.**
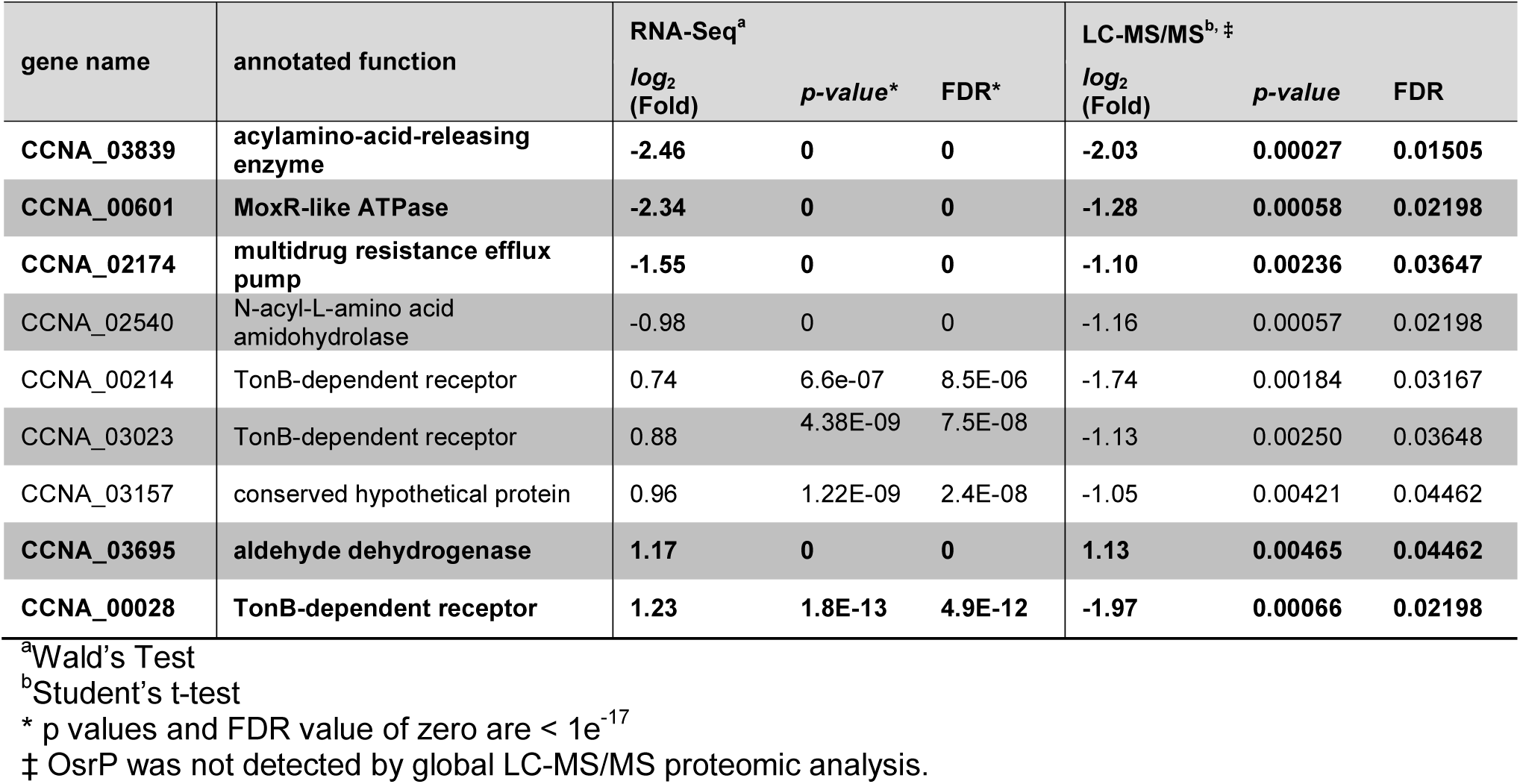
Proteins with significant differences in steady-state levels between *gsrN*^++^ and wild type (*gsrN*^++^ / WT) during hyperosmotic stress, with associated transcript changes.

**FIG 3.**
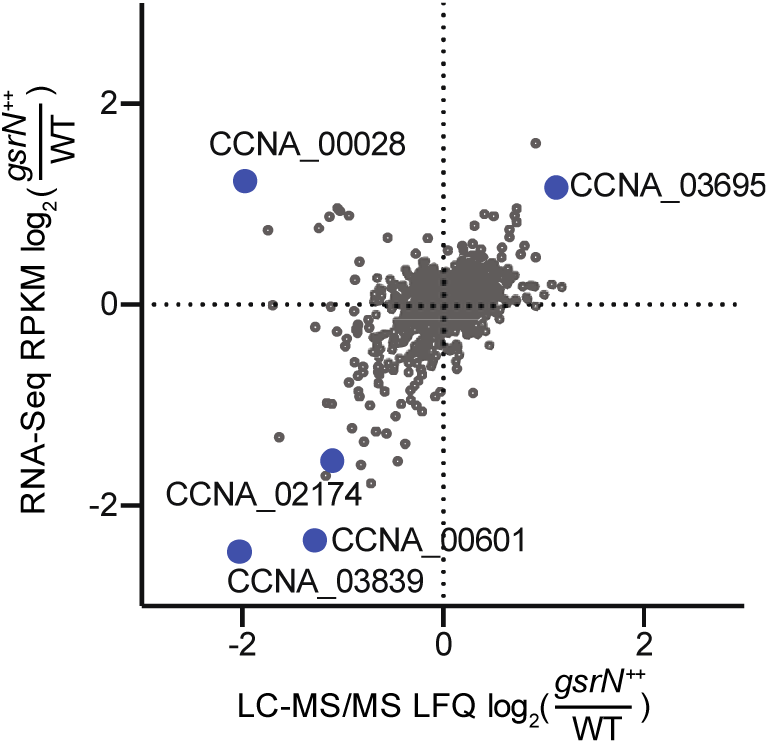
*gsrN*-regulated genes under hyperosmotic conditions. Transcriptomic and proteomic analysis of *gsrN*^++^ and wild type (WT) strains after sucrose-induced hyperosmotic stress. Relative reads per kilobase per million (RPKM) for transcriptomics and label-free quantitation (LFQ) for proteomics are plotted. Only genes detected in both analyses are plotted. Blue points indicate genes for which transcript and protein levels differed significantly between *gsrN*^++^ and WT. Significant differential regulation cutoff was *log_2_*(fold) > 1.0 and FDR *p-value* < 0.05 for both transcript and protein based on Wald’s Test and Student’s t-test, respectively. RNA-Seq data set comprises of 3 wild-type unstressed, 3 wild-type stressed, 4 *gsrN*^++^ unstressed, and 4 *gsrN*^++^ stressed conditions. LC-MS/MS data set comprises the same number of samples for each respective strain and treatment.

### Comparative RNA-seq analysis uncovers candidate targets of GsrN under hyperosmotic stress

To delineate the roles of *sigT* and *gsrN* in stress survival, we compared genes that are differentially regulated between Δ*sigT, gsrN*^++^ and wild type strains subjected to hyperosmotic stress. More explicitly, we sought to test the model that GsrN functions as a direct post-transcriptional regulator of mRNAs that are dependent on σ^T^-transcription. Since transcription of *gsrN* is directly activated by σ^T^, we expected (in this model) that the set of transcripts modulated by *gsrN* overexpression should exhibit some overlap with the set of transcripts that changes upon *sigT* deletion. Indeed, we observed 20 genes with congruent patterns of regulation in these two datasets (**Fig. 4A and Table 3**).

**Table 3.** RNA-seq analysis of Δ*sigT* and *gsrN*^++^ during hyperosmotic stress.

| gene name | annotated function | $gsrN^{++}/WT$ | | | $\Delta sigT/WT$ | | |
| --- | --- | --- | --- | --- | --- | --- | --- |
| | | $\log_2$ (Fold) | $p$-value<sup>1</sup> | FDR<sup>1</sup> | $\log_2$ (Fold) | $p$-value<sup>1</sup> | FDR<sup>1</sup> |
| *gsrN* | cell cycle regulated sRNA *gsrN* | 3.14 | 0 | 0 | -6.27 | 7.98E-14 | 3.33E-12 |
| *CCNA_00882* | hypothetical protein | 2.75 | 0 | 0 | -5.82 | 0 | 0 |
| *CCNA_03694* | AcoR-family transcriptional regulator | 1.37 | 0 | 0 | -1.64 | 0 | 0 |
| ***CCNA_00028\**** | TonB-dependent receptor | 1.23 | 1.82E-13 | 4.90E-12 | -1.32 | 2.85E-14 | 1.27E-12 |
| ***CCNA_03695\**** | aldehyde dehydrogenase | 1.17 | 0 | 0 | -1.40 | 1.10E-05 | 1.52E-04 |
| *CCNA_03889* | conserved hypothetical protein | 1.10 | 1.55E-04 | 1.25E-03 | -1.30 | 1.92E-12 | 7.13E-11 |
| *CCNA_00709* | hypothetical protein | 1.07 | 5.67E-04 | 3.89E-03 | -6.22 | 3.90E-06 | 5.97E-05 |
| *CCNA_01089* | conserved hypothetical protein | -1.00 | 1.11E-16 | 4.12E-15 | 1.18 | 5.44E-13 | 2.10E-11 |
| *CCNA_02051* | imidazolonepropionase related amidohydrolase | -1.22 | 0 | 0 | 1.11 | 3.09E-09 | 7.70E-08 |
| *CCNA_01653* | cyclophilin-type peptidylprolyl cis-trans isomerase | -1.23 | 0 | 0 | 1.39 | 0 | 0 |
| *CCNA_00243* | hypothetical protein | -1.33 | 0 | 0 | 1.25 | 3.13E-11 | 9.84E-10 |
| *CCNA_03082* | hypothetical protein | -1.34 | 0 | 0 | 1.24 | 2.82E-12 | 1.03E-10 |
| *CCNA_01100* | acylamino-acid-releasing enzyme | -1.55 | 0 | 0 | 1.16 | 9.52E-10 | 2.56E-08 |
| ***CCNA_02174\**** | multidrug resistance efflux pump | -1.55 | 0 | 0 | 2.32 | 0 | 0 |
| *CCNA_02935* | methyl-accepting chemotaxis protein | -1.66 | 0 | 0 | 2.12 | 0 | 0 |
| *CCNA_02172* | ABC-type transporter, permease component | -1.72 | 0 | 0 | 2.37 | 0 | 0 |
| *CCNA_02173* | ABC transporter ATP-binding protein | -1.79 | 0 | 0 | 2.29 | 0 | 0 |
| *CCNA_01247* | CESA-like glycosyltransferase | -2.00 | 0 | 0 | -1.53 | 6.29E-10 | 1.72E-08 |
| *CCNA_02050* | imidazolonepropionase related amidohydrolase | -2.10 | 0 | 0 | 1.29 | 2.91E-08 | 6.51E-07 |
| *CCNA_03687* | carbonic anhydrase | -2.24 | 0 | 0 | 1.99 | 0 | 0 |

Genes also identified in Fig. 3 <sup>1</sup> p values and FDR value of zero are < 1e-17

**FIG 4.**
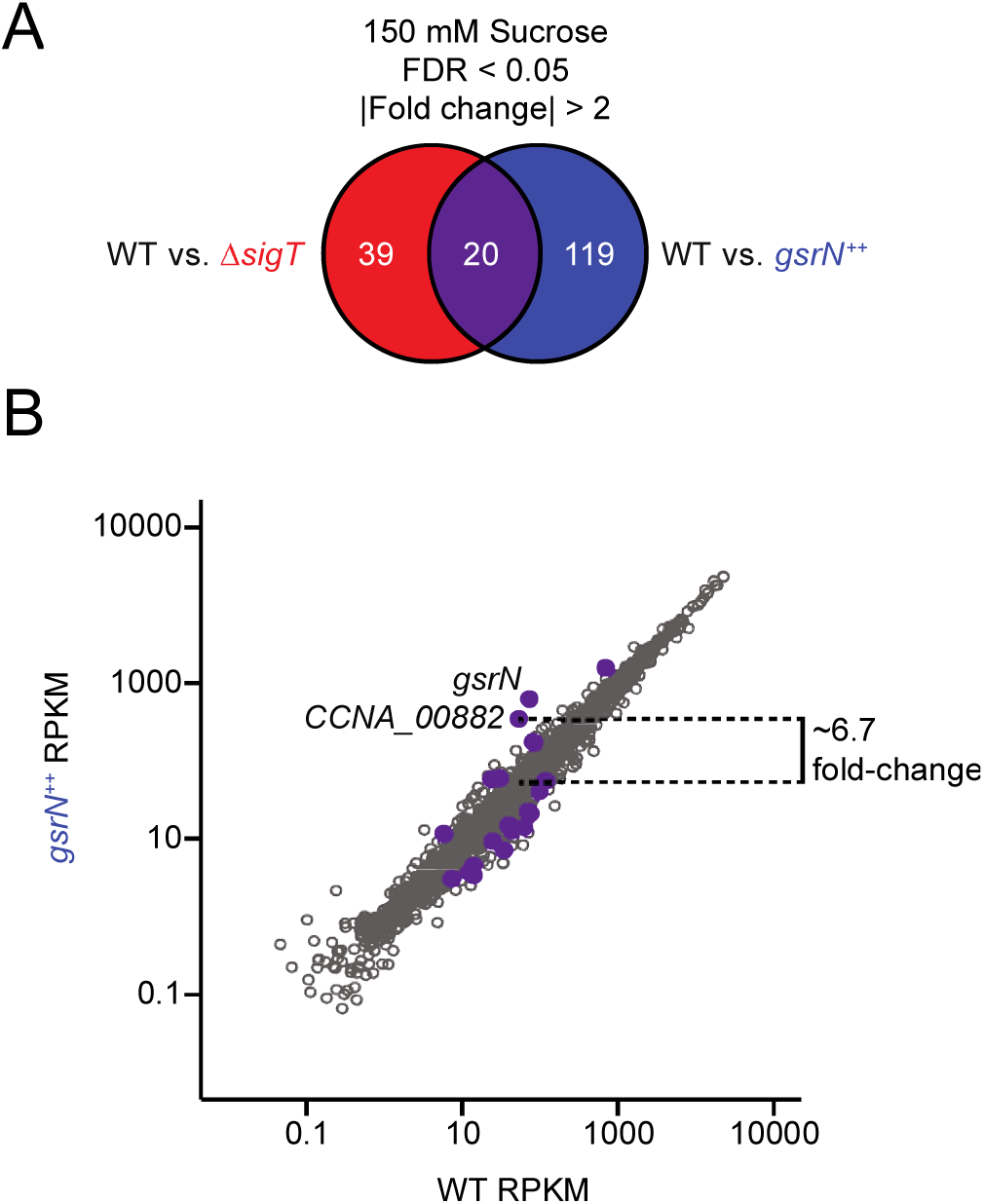
Comparative RNA-seq analysis uncovers candidate targets of GsrN under hyperosmotic stress. (A) RNA-Seq Venn summary of genes differentially regulated in Δ*sigT* (red) and *gsrN*^++^ (blue) during sucrose-induced hyperosmotic stress; the shared gene set is highlighted in purple. Significant differential regulation cutoff was *log_2_*(fold) > 1.0 and FDR *p-value* < 0.05 for both comparisons. (B) Measured transcript abundance, reads per kilobase per million (RPKM), in WT and *gsrN*^++^ samples subjected to sucrose-induced hyperosmotic stress. Purple points represent genes identified in (A). Dotted line highlights the 2.7 *log_2_*(fold) change of *CCNA_00882* (*osrP*) between *gsrN*^++^ and WT.

In this set of 20 genes, we identified six candidate GsrN target genes whose transcript levels were significantly lower in Δ*sigT* and higher in *gsrN*^++^ under hyperosmotic stress (**Fig. 4B**). We predicted strong σ^T^-binding sites in the promoters of five of these candidates: *CCNA_00882*, *CCNA_00709*, *CCNA_03889*, and *CCNA_03694-CCNA_03595* (**Table S1**). Of these candidate direct regulatory targets of GsrN, *CCNA_00882* showed the highest change (≈7 fold) upon overexpression of *gsrN* (**Fig. 4B**). Moreover, steady-state *CCNA_00882* transcript levels were significantly higher during osmotic stress in wild type cultures (≈6 fold) and in GsrN overexpression (*gsrN*^++^) cultures (≈12.5 fold). Thus, we named *CCNA_00882*, *osrP*, <u>o</u>smotic <u>s</u>tress <u>r</u>egulated <u>p</u>rotein. We note that *osrP* mRNA was previously identified as an RNA that co-elutes with GsrN in an affinity pull-down experiment (11). Considering the presence of a σ^T^-binding motif in the *osrP* promoter, its regulation by GsrN in our transcriptomic datasets, and the fact that it co-purifies with GsrN, we postulated that o*srP* is a direct target of GsrN.

### *osrP* is regulated by σ^T^, induced under hyperosmotic stress, and interacts with GsrN via its 5’ leader sequence

*osrP* is annotated as a 332-residue hypothetical protein that is largely restricted to the genus *Caulobacter*, based on a BLAST search (22) of the GenBank non-redundant database. However, the primary structure of *osrP* shares some features with annotated open reading frames across genera in the family Caulobacteraceae including *Phenylobacterium*, *Asticcacaulis*, and *Brevundimonas*. OsrP contains a signal peptide at its amino terminus with a Type I signal peptidase cleavage site, as predicted by SignalP (23). A conserved glycine zipper motif (Pfam05433) comprised of two hydrophobic helices is located between residues 224 and 268. Based on these sequence features, we predict that OsrP is a periplasmic protein **(Fig. S3)**.

To better understand the regulation of *osrP* by σ^T^ and GsrN, we identified its transcription start site (TSS) by 5’ rapid amplification of cDNA ends (5’ RACE). We mapped the *osrP* TSS to nucleotide 962935 in the *C. crescentus* genome (Genbank accession NC_011916); a near-consensus σ^T^ binding site is positioned at −35 and −10 relative to the *osrP* TSS (**Fig. 5A and Table S1**). To assess transcriptional regulation of *osrP*, we generated a fusion of the *osrP* promoter to a promoterless *lacZ* (**Fig. 5B**). β-galactosidase activities in wild-type and Δ*gsrN* strains harboring this reporter plasmid were comparable under untreated conditions, and transcription was activated in both of these genetic backgrounds upon addition of 150 mM sucrose to induce hyperosmotic stress. In a Δ*sigT* strain, we observed basal β-galactosidase activity in untreated conditions, and activity was not induced upon addition of 150 mM sucrose (**Fig. 5B**). We conclude that transcription of *osrP* depends on *sigT* and is independent of *gsrN*.

**FIG 5.**
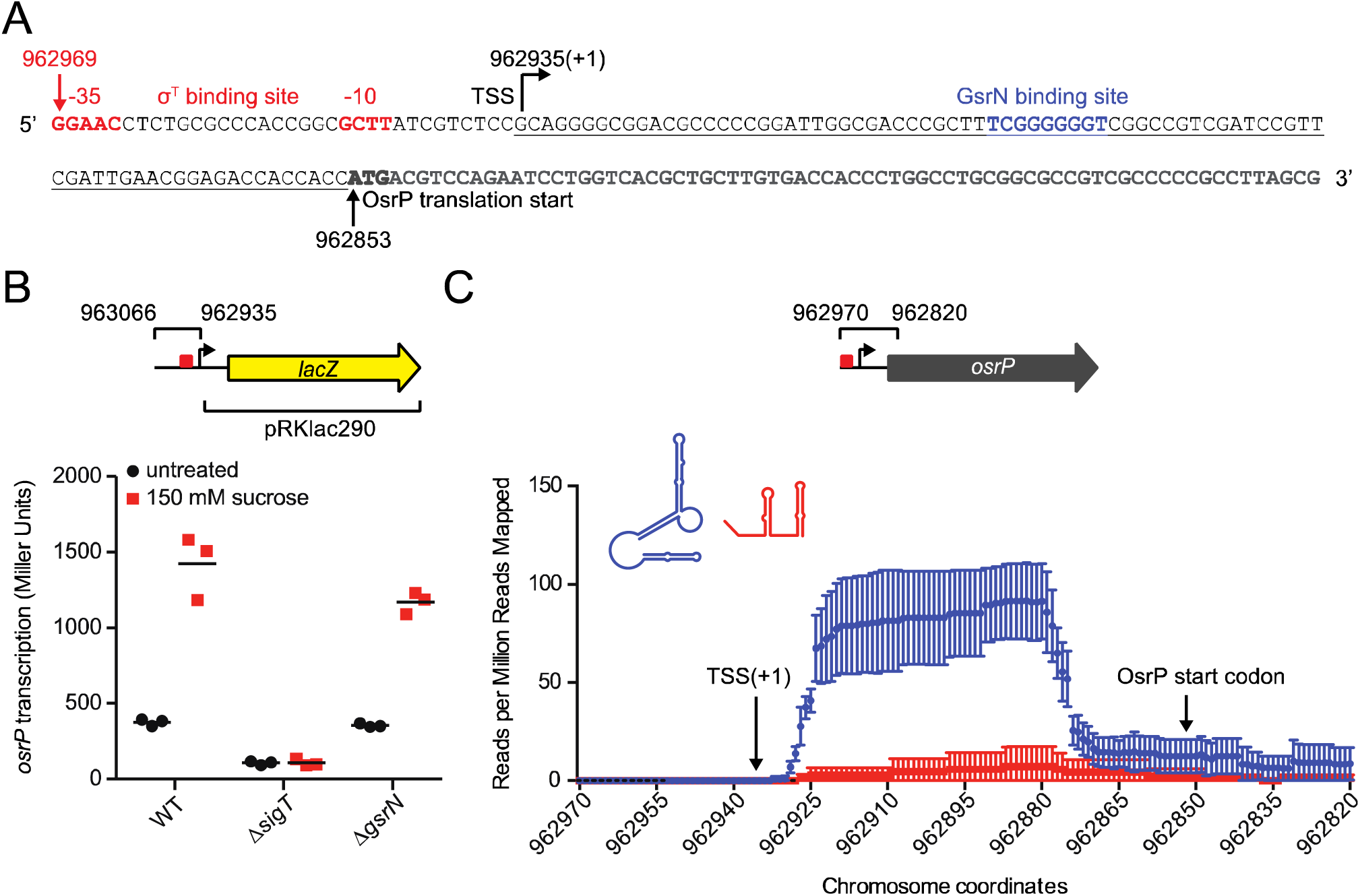
*osrP* is transcribed by σ^T^ and is upregulated during hyperosmotic stress. (A) *osrP* promoter and leader sequence. Bent arrow indicates the location of the transcription start site (TSS) mapped by 5’ RACE. The proposed σ^T^-binding site at −10 and −35 is in red. The proposed GsrN binding site from (11) is in blue. Black arrow and bolded nucleotides indicate the annotated translation start site of *osrP*. The 5’ leader of *osrP* mRNA is underlined. Numbers correspond to nucleotide positions in genome accession NC_011916 (B) β-galactosidase activity assay (in Miller Units) of the pRKLac290-*osrP* transcriptional reporter plasmid in wild type (WT), Δ*sigT*, and Δ*gsrN* backgrounds; a schematic of the reporter plasmid marking the cloned region of the *osrP* promoter with the σ^T^-binding site (red) is pictured above. Activities in log phase cultures without or with 150 mM sucrose (3 hour treatment) are in black circles and red squares, respectively. Horizontal bars mark the mean of three independent biological replicates. (C) mRNA that co-purified with *gsrN(37)::PP7hp* (aptamer-tagged GsrN; blue) and *PP7hp::gsrN-3’* (negative control; red) quantified as fractional reads mapped to the leader region of *osrP*. Read density in each dataset represents read coverage at each nucleotide divided by the number of million reads mapped in that data set. Data represent mean ±SD of three replicate *gsrN(37)::PP7hp* and two replicate *PP7hp::gsrN-3’* purifications.

We previously affinity purified GsrN tagged with a PP7 RNA hairpin aptamer (GsrN(37)::PP7hp) from *Caulobacter* cell lysate, and quantified RNAs that co-purified with GsrN by RNA-seq (11). For this present study, we have re-analyzed our published dataset (NCBI GEO accession number GSE106171) with the goal of identifying reads that map to *osrP* mRNA. We observed significant enrichment of reads that map to the extended 5’ leader sequence of *osrP*, which is comprised of approximately 80 nucleotides between the TSS and the annotated start codon. No enrichment of the *osrP* leader is observed with the PP7hp::GsrN-3’ negative control (**Fig. 5C**). IntaRNA analysis (24) of this co-purifying region predicted strong base-pairing between the 5’ C-rich loop of GsrN and the 5’ leader of *osrP* (**Fig. 5A**). From these data, we conclude that GsrN interacts with the 5’ untranslated leader of *osrP* mRNA.

### GsrN activates OsrP expression at the post-transcriptional level

To test the functional significance of the proposed GsrN binding site in the 5’ leader of *osrP* mRNA, we constructed an *osrP* transcriptional-translational (TT) reporter plasmid. The reporter contains the *osrP* promoter, the 5’ untranslated region (5’ UTR), and the first 7 codons of *osrP* fused to 5’ end of *lacZ* lacking a start codon (**Fig. 6A**). The RNA-fold (25) structure of the 5’ UTR and the nucleotides encoding the first 7 amino acids of *osrP* predicts that the majority of the GsrN-binding site is sequestered in a base-paired region (26) (**Fig. 6B**). Under unstressed conditions, activity from the TT reporter is low in wild-type cells, but reduced by 2 fold in Δ*sigT* and Δ*gsrN* backgrounds. Overexpression of *gsrN* (*gsrN*^++^) enhances expression 8 fold compared to wild type. This enhancement of *osrP* expression requires *sigT*, as overexpression of *gsrN* from the σ ^70^ P1 promoter in a Δ*sigT* background does not induce expression from the *osrP* TT reporter **(Fig. 6C)**. This result is consistent with data presented in **Fig. 5B** showing the *osrP* transcription requires σ^T^ and supports a model in which GsrN regulates OsrP protein expression at the post-transcriptional level. Lastly, our measurements of *osrP* TT reporter activity under hyperosmotic stress conditions showed similar relative regulatory trends across the assayed genetic backgrounds, though baseline expression is higher (**Fig. 6C**).

**FIG 6.**
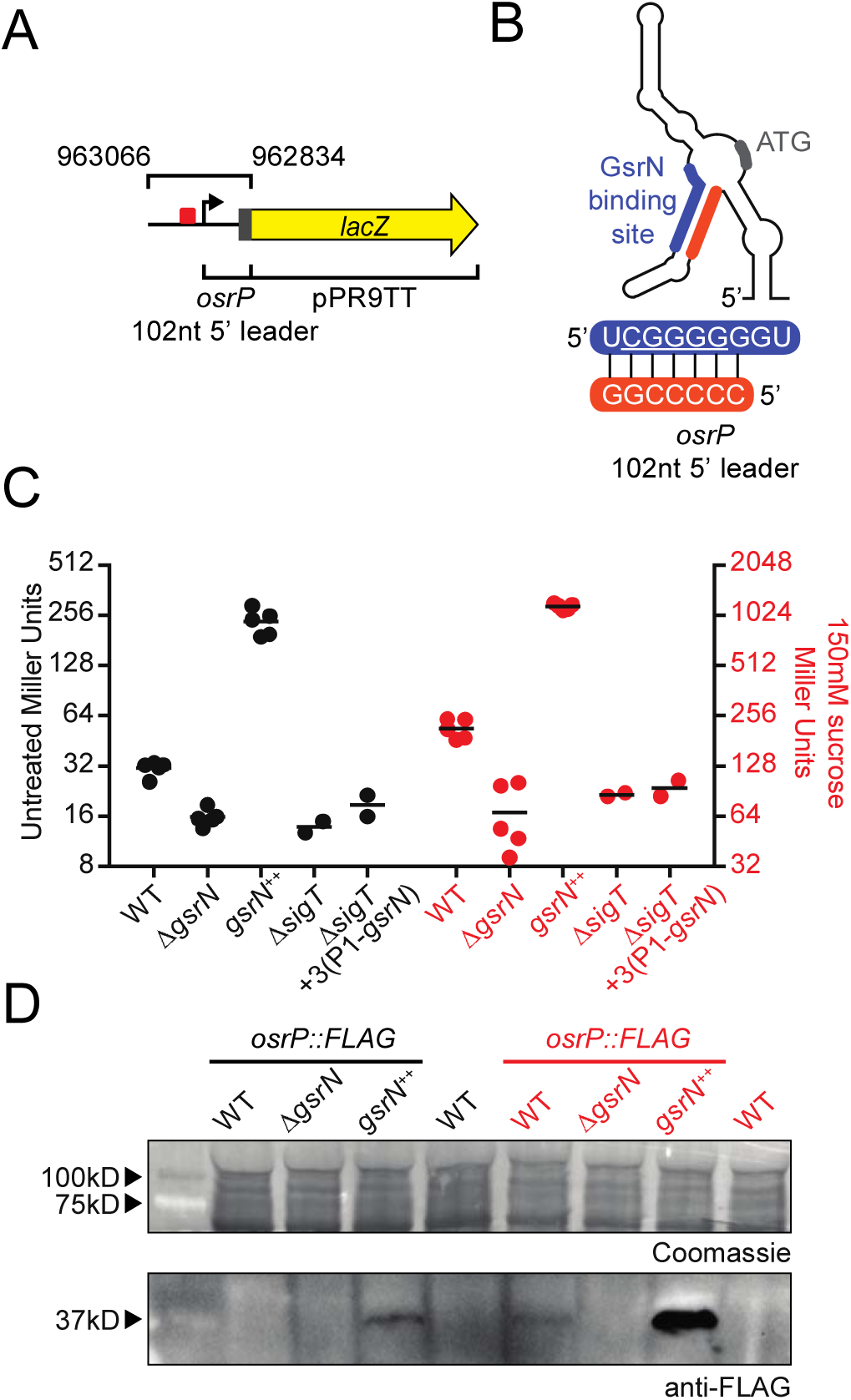
GsrN activates the expression of OsrP at the post-transcriptional level. (A) Schematic of pPR9TT-*osrP* transcription plus translation (TT) reporter plasmid. The nucleotide positions of the *osrP* genome region (upstream region, 5’ UTR, and nucleotides encoding the first 7 amino acids) fused to *lacZ* are indicated on the upper bracket. 5’ leader of *osrP* (5’ UTR and nucleotides encoding the first 7 amino acids) is marked with the bottom bracket. (B) Predicted secondary structure (26) of 5’ leader of *osrP*. A proposed GsrN binding site is highlighted in blue (see also Fig 5). Base-paired region complementary to the proposed GsrN binding site is highlighted in orange. Start codon is highlighted in grey. Sequence below shows the interaction between the predicted GsrN-binding site and the complementary base-paired region within the 5’ leader of *osrP*. (C) β-galactosidase activity assay (in Miller Units) from the pPR9TT-*osrP* TT reporter plasmid. Left black axis represents reporter activity in untreated cultures. Right red axis represents reporter activity in cultures treated with 150 mM sucrose for 3 hours. Data and mean represent at least two biological replicates. (D) Western analysis of total protein from WT, Δ*gsrN*, and *gsrN*++ strains containing *osrP::FLAG*, Untreated and hyperosmotic treated (150 mM sucrose) cultures are black and red, respectively. After transfer, the top portion of the membrane was Coomassie-stained as a loading control (top panel). The bottom portion of the membrane was blotted with anti-FLAG antibodies (lower panel). Blot is overexposed to reveal weaker bands. As a result, the OsrP::FLAG signal in the *gsrN*^++^ treated lane is saturated. Arrows on the left indicate protein size markers. Blot and stained membrane are representative of duplicate experiments.

To directly measure OsrP protein levels, we inserted a C-terminal FLAG tag at the native *osrP* locus on the *Caulobacter* chromosome. OsrP::FLAG is expressed at low levels in exponentially growing wild type cultures, and was difficult to detect by Western blot. Steady-state levels of OsrP::FLAG were higher in cultures overexpressing GsrN (*gsrN*^++^) (**Fig. 6D**). Hyperosmotic stress (150 mM sucrose) induced production of OsrP::FLAG in wild-type cells. OsrP::FLAG detection required *gsrN*: there was no detectable OsrP::FLAG in Δ*gsrN* cells. Consistent with our *osrP* reporter data, we observed a large increase in OsrP::FLAG levels in *gsrN*^++^ relative to wild type during hyperosmotic stress (**Fig. 6D**). Again, these data support a model in which GsrN post-transcriptionally activates OsrP protein expression.

### Assessing the role of the GsrN C-rich recognition loop in activation of OsrP expression

It is established that the C-rich target recognition loop is a functionally-important feature of GsrN structure that directly activates KatG catalase/peroxidase expression through a base-paring interaction with the 5’ leader of *katG* mRNA (11). To test the role of this recognition loop in the activation of OsrP protein expression, we constructed a TT *osrP* reporter containing reverse-swapped (RS) mutations in the 5’ leader of *osrP* (*osrP*-RS1). These mutations were expected to disrupt predicted base-pairing interactions with wild-type GsrN and restore base pairing interactions with the complementary GsrN(RS) recognition loop mutant (see **Fig. S4**).

It is important to note that the RS1 mutations disrupt predicted secondary structure **(Fig. S4A)** and calculated stability of the *osrP* leader (mFold −46.6 kcal/mol for wild-type compared to −39.8 kcal/mol for *osrP*-RS1 (26)). The *osrP*-RS1 reporter had substantially higher basal reporter activity than the wild-type *osrP* TT reporter (Fig. S4B and S4F). Thus the RS1 mutations alone disrupt the *osrP* leader structure and regulation of OsrP expression, making it difficult to infer a regulatory role for base pairing with GsrN. Deletion of *gsrN* had no effect on *osrP*-RS1 activity, as expected if base pairing is disrupted by the RS1 mutations. However, overexpression of *gsrN* unexpectedly activated expression from the *osrP*-RS1 reporter **(Fig. S4B)**. In the opposite experiment, in which we engineered complementary RS mutations into the GsrN recognition loop, the activity from the wild-type *osrP* TT reporter was diminished **(Fig. S4C)**. In the case of direct base pairing, we would expect RS reporter activity to be restored when complementary RS mutations are present in both GsrN and the *osrP* leader. However, the high basal activity of *osrP*-RS1 in the strain bearing both these mutations poses challenges in interpretation of these data.

Because the RS1 mutations disrupt structure, stability and regulation of the *osrP* leader, we sought to compensate for base pairing disruptions arising in *osrP*-RS1 by introducing compensatory base changes on the opposing arm of the RNA stem in the *osrP* leader. These mutations were predicted to restore the secondary structure and the stability of the leader; we termed this the *osrP*-RS2 reporter **(Fig. S4D)**. We failed to detect any activity from the *osrP*-RS2 reporter in any strain, including those expressing *gsrN(RS)* (**see Fig. S4E-F**). The lack of activity could be due to the lower levels of GsrN(RS) versus wild-type GsrN (11) or it may be the case that the RS2 mutations ablate an essential regulatory feature of the *osrP* leader. Based on these reporter data, we conclude that mutation of the predicted GsrN target site in the 5’ leader of *osrP* derepresses expression of this protein. From these experiments, we are not able to define the base pairing interaction between the GsrN C-rich loop and the 5’ leader of *osrP* mRNA that activates OsrP expression.

### *osrP* does not affect hyperosmotic stress survival

As *osrP* expression is under strong positive control of σ^T^ and GsrN during hyperosmotic stress, we tested the possibility that *osrP* contributes to stress survival. Deletion of *osrP* (Δ*osrP*) did not affect survival of *C. crescentus* under hyperosmotic conditions **(Fig. S1D)**. Expression of *osrP* from a xylose-inducible expression plasmid (*osrP*^++^) also had no effect on hyperosmotic stress survival **(Fig. S1D)**. We further tested the role of *osrP* on osmotic stress survival in Δ*gsrN* and *gsrN*^++^ backgrounds. *osrP* is not required for the protective effect conferred by *gsrN*^++^ (see ∆*orsP gsrN*^++^ in **Fig. S1D**) and overexpression of *osrP* does not rescue strains lacking *gsrN* (see ∆*gsrN osrP*^++^ in **Fig. S1D**). From these data, we conclude that *osrP* is not the sole contributor to hyperosmotic stress survival under the assayed conditions.

## Discussion

Microbes employ regulatory systems that function to mitigate the effects of osmotic stress (27). The freshwater oligotroph *Caulobacter crescentus* activates the general stress response (GSR) sigma factor, σ^T^, during hyperosmotic stress. This, in turn, activates transcription of a large set of genes **(Table S3)** including the sRNA, GsrN (11, 21). Deleting either *sigT* or *gsrN* results in reduced viability under sucrose-induced hyperosmotic stress **(Fig. 1C and Fig. 2C)**. Unlike oxidative stress, in which *gsrN* expression alone is sufficient to protect cells (11), *gsrN*-dependent protection of *Caulobacter* during hyperosmotic stress requires that the *sigT* gene remain intact **(Fig. 2C)**.

Transcriptomic analysis of a Δ*sigT* strain provided a comprehensive view of the σ^T^ hyperosmotic stress regulon, while transcriptomic and proteomic analysis of a *gsrN* overexpression strain (*gsrN*^++^) revealed a set of transcripts and proteins that are under post-transcriptional control of GsrN during hyperosmotic stress **(Fig. 3 and Table 2)**. Comparative analyses of these datasets provided evidence for multi-output feedforward loops (FFL) involving σ^T^ and GsrN. One such coherent FFL involves the uncharacterized glycine-zipper protein, OsrP. Specifically, transcription of *osrP* is activated by σ^T^ **(Fig. 5B)**, likely via direct binding to the canonical σ^T^ binding site in its promoter **(Fig. 5A and Table S3)**. OsrP protein expression is activated at the post-transcriptional level by GsrN **(Fig. 6C-D)**, to form a coherent FFL. Both *sigT* and *gsrN* are required for OsrP protein expression; thus, both regulators comprise an AND gate that regulates *osrP* expression.

### A coherent feedforward loop controls a *Caulobacter* gene expression response during hyperosmotic stress

Feedforward loops (FFL) are common regulatory motifs in microbial gene expression networks. In their simplest form, FFLs are comprised of three genetic components: two regulators and an output gene. The primary regulator functions to activate both the secondary regulator and the output gene; the secondary regulator functions to activate expression of the output gene (28). In the case of *osrP*, σ^T^ is the primary regulator that activates *osrP* and *gsrN* transcription; GsrN interacts with *osrP* mRNA to activate OsrP protein expression at the post-transcriptional level **(Fig. 7)**.

**FIG 7.**
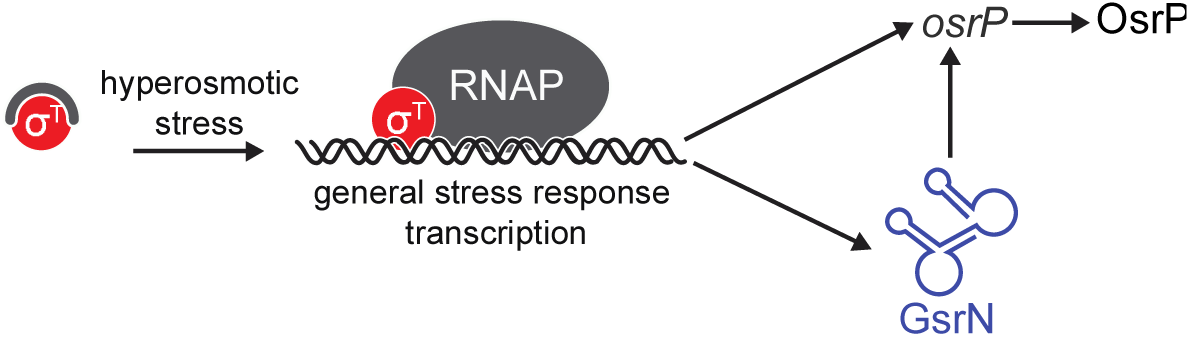
Coherent feedforward regulation during hyperosmotic stress in *C. crescentus*. σ^T^ is de-repressed upon exposure to hyperosmotic stress, and binds to core RNA polymerase (RNAP). σ^T^ subsequently activates transcription of a set of genes (see Table S3), including the sRNA, GsrN. GsrN accumulates in the cell and functions to either activate or repress expression of genes at the post-transcriptional level (see Table S4). *osrP* is among a set of genes in the GSR hyperosmotic stress regulon that are upregulated by σ^T^ at the transcriptional level and also upregulated by GsrN at the post-transcriptional level, i.e. coherent feedforward regulation.

There are several examples of sRNAs that are part of FFL motifs in bacteria (29–31). Activation of *osrP* expression by σ^T^ and GsrN in *Caulobacter* is perhaps most similar to the regulation of *ricI* by σ^S^ and the sRNA RprA in *Salmonella* (30); RicI functions as an inhibitor of plasmid transfer. In this instance, the primary and secondary regulators are swapped: RprA acts as a primary activator of *rpoS* and *ricI* expression. RprA itself is activated by the Rcs system (32), which responds to envelope stress. However, *rpoS* expression is controlled by multiple environmental signals and does not require *rprA* to transcribe *ricI*. Thus, *ricI* can be transcribed in the absence of envelope stress, but both RprA and σ^S^ are required for RicI protein expression. Thus, RprA and σ^S^ function as a FFL AND gate that ensures RicI expression only occurs upon Rcs activation by envelope damage.

Unlike *ricI*, the coherent FFL controlling *osrP* is activated by σ^T^ alone. In wild-type *C. crescentus*, we observe basal σ^T^-dependent gene expression in the absence of any apparent stress **(Fig. 5B and Fig 6C)**. During hyperosmotic stress, *osrP* expression measured from the *osrP i)* transcription and *ii)* transcription plus translation (TT) reporters are incongruent: *osrP* transcription increases 2-fold during hyperosmotic stress while TT reporter activity increases 6-fold within an equivalent time window. The difference in fold change between the two reporters is likely due to the positive regulatory effects of GsrN, the levels of which increase 3-fold during hyperosmotic stress (11).

During persistent stress conditions, such as hyperosmotic stress and stationary phase, we have observed that the stable 5’ isoform of GsrN accumulates to higher levels than full-length GsrN (11). Accumulation of the 5’ GsrN isoform could act as signal within the *sigT*-regulon to mount a specific response to persistent stress, such as hyperosmotic shock. This model is consistent with AND-type coherent FFLs, which result in delayed activation of the output gene. Expression delay arises from the lag between the production of the secondary regulator and the threshold necessary for the secondary regulator to act upon the output gene (28). In the case of *osrP*, levels of GsrN may set the threshold for OsrP protein production. Accumulation of the 5’ GsrN isoform through prolonged σ^T^-activity could amplify the expression of *osrP* over other σ^T^-regulated genes in particular stress regimes. Although we conclusively demonstrate only one GSR coherent feedforward loop (during hyperosmotic stress) in this study, our transcriptomic and proteomic data show that several genes in the GSR regulon may be subject to similar regulation.

### Functional analysis of the uncharacterized glycine zipper protein, OsrP

Bioinformatic analysis of OsrP predicts two notable features in its sequence: a signal peptide at its amino terminus with a Type I signal peptidase cleavage site and a conserved glycine zipper motif (Pfam05433) **(Fig. S3)**. From this analysis, we predict that OsrP is in the periplasm of *C. crescentus*. The primary structure of the glycine zipper motif suggests a possible interaction with the cell membrane. Extended glycine zipper motifs can oligomerize and form pores within membranes (19). One notable example is the secreted VacA toxin of *H. pylori* that forms a hexameric anion selective channel in host cells (33).

In considering its primary structure, predicted localization, and regulation, it seemed possible that *osrP* could help alleviate osmotic stress in *C. crescentus.* However, deletion of *osrP* did not result in any obvious viability defect during sucrose-induced hyperosmotic stress. There are several genes co-regulated by σ^T^ and GsrN, and it may be the case that additional genes are required to mitigate the hyperosmotic stress conditions that we have tested. Prolonged exposure to other osmotic stresses and/or different concentrations of osmolytes – including a range of ions – could provide insight into function of *osrP* activation by the GSR.

In a recent study of a diverse set of bacterial species, including *C. crescentus*, growth of transposon mutant libraries was characterized under multiple environmental conditions (34). In *C. crescentus*, *osrP* disruption resulted in a consistent disadvantage in growth in the presence of sodium perchlorate (fitness = −1.3, t score =-7.2). Sodium perchlorate is an anionic oxidizing agent. It remains uncertain how the oxidative, osmotic (or other) effects of sodium perchlorate in the medium affect fitness of strains harboring transposon disruptions of *osrP*, but this result provides an additional assay condition for future functional studies of *osrP*. Considering this sodium perchlorate result, we assayed survival of the *osrP* deletion strain under peroxide stress. The *osrP* deletion strain had no survival defect in the face of 200 μM hydrogen peroxide exposure for 1 hour, while a *sigT* deletion strain had an expected ~2–3 log defect as previously reported (11). Studies to uncover conditions under which an *osrP* mutant has a growth or survival phenotype is ongoing.

### On additional GsrN regulatory targets

Proteomic analysis of the *gsrN*^++^ strain showed different sets of regulated genes between untreated and hyperosmotic stress conditions. In untreated cultures of *gsrN*^++^, all proteins with significant differential expression were under negative control of GsrN; there was no overlap with the differentially expressed proteins we observed in stress-treated cultures. Although GsrN may not directly control expression of all differentially regulated proteins in this dataset, the effect we observe upon *gsrN* overexpression in the absence of stress points to a role for GsrN during normal growth **(Table 1)**. Notably, the cell-cycle phosphotransferase, ChpT, is significantly downregulated upon *gsrN* overexpression. ChpT is an essential protein is required for phosphorylation of the essential cell cycle master regulator, CtrA (35). Given the established connection between levels of CtrA and σ^T^ during nutrient limitation (36), it is conceivable that GsrN regulates the core cell cycle control system of *C. crescentus* under certain conditions.

Under hyperosmotic conditions, we observed only a few cases of proteins that differ significantly in steady-state levels between the *gsrN*^++^ and wild type strains. Besides *osrP*, genes regulated by GsrN during hyperosmotic stress identified in this study were not identified in our previous study of mRNAs that co-purify with GsrN-PP7 (11). However, affinity-purification of GsrN-PP7 and its co-eluting RNAs was performed in unstressed conditions. Identification of co-eluting mRNAs using the PP7 aptamer pull-down approach is biased toward highly expressed RNAs; *osrP* mRNA was the highest expressed target during hyperosmotic stress (Figure 4). Future pull-down experiments conducted under a variety of stress conditions may define new GsrN mRNA targets that were missed due to low steady-state levels under non-inducing conditions.

Among the proteins negatively regulated by GsrN are three TonB-dependent receptors of unknown function (*CCNA_00028*, *CCNA_00214*, and *CCNA_3023*) and a predicted efflux complex (*CCNA_02172-74*) **(Tables 2 and 3)**. GsrN may therefore have a functional role that is similar to the sRNAs, MicA and RybB, which are transcribed during envelope stress by σ^E^ in *E. coli* and repress outer membrane proteins (OMP) to mitigate accumulation of unfolded OMPs (37). Genes activated at the transcript level by GsrN during hyperosmotic stress include several with predicted σ^T^-binding sites in their promoters (*CCNA_00709*, *CCNA_03889*, and *CCNA_03694-CCNA_03595*). These may provide additional cases of coherent FFLs. As discussed previously, expression of these genes may be sensitive to GsrN accumulation during prolonged stress. CCNA_00709 – a predicted small, two-pass membrane protein – and CCNA_03694 – a transcription factor – are attractive targets to investigate in future studies on the mechanism by which GsrN determines cell survival during hyperosmotic stress.

## Materials and Methods

All *C. crescentus* experiments were conducted using strain CB15 (38) and derivatives thereof.

### Growth of *E. coli* and *C. crescentus*

*C. crescentus* was cultivated on peptone-yeast extract (PYE)-agar (0.2% peptone, 0.1% yeast extract, 1.5% agar, 1 mM MgSO_4_, 0.5 mM CaCl_2_) (39) at 30°C. Antibiotics were used at the following concentrations on this solid medium: kanamycin 25 μg/ml, tetracycline 2 μg/ml, nalidixic acid 20 μg/ml, and chloramphenicol 2 μg/ml. For liquid culture, *C. crescentus* was cultivated in either PYE or in M2X defined medium (39). PYE liquid: 0.2%(w/v) peptone, 0.1%(w/v) yeast extract, 1 mM MgSO_4_, and 0.5 mM CaCl_2_, autoclaved before use. M2X defined medium: 0.15% (w/v) xylose, 0.5 mM CaCl_2_, 0.5 mM MgSO_4_, 0.01 mM Fe Chelate, and 1x M2 salts, filtered with a 0.22 micron bottle top filter. One liter of 20x M2 stock was prepared by mixing 17.4 g Na_2_HPO_4_, 10.6 KH_2_PO_4_, and 10 g NH_4_Cl. Antibiotics were used at the following concentrations in liquid medium: kanamycin 5 μg/ml, tetracycline 1 μg/ml, and chloramphenicol 2 μg/ml. For cultivation of *E. coli* in liquid medium, we used lysogeny broth (LB). Antibiotics were used at the following concentrations: kanamycin 50 μg/ml, tetracycline 12 μg/ml, and chloramphenicol 20 μg/ml.

### Plasmid transformation into *C. crescentus*

Plasmids were conjugated into CB15 (39) using the *E. coli* helper strain FC3 (40) **(see Table S1)**. Conjugations were performed by mixing the donor *E. coli* strain, FC3, and the CB15 recipient strain in a 1:1:5 ratio. Mixed cells were pelleted for 2 min at 15,000xg, resuspended in 100 μL, and spotted on a nonselective PYE-agar plate for 12–24 hr. Exconjugants containing the desired plasmid were selected on PYE agar containing the plasmid-specified antibiotic for selection and nalidixic acid (20 μg/ml) to counter-select against both *E. coli* strains (helper and plasmid donor). Plasmids pMT552 and pMT680 integrate into the *vanA* and *xylX* locus respectively. pMT680 carries a chloramphenicol resistance marker gene (*cat*) and pMT552 carries a kanamycin resistance marker gene (*npt1*) (41). pNPTS138 integration occurs at a chromosomal site homologous to the insertion sequence.

### Chromosomal deletion and allele replacement in *C. crescentus*

To generate the in-frame deletion and C-terminal FLAG-tagged *osrP* (*CCNA_00882*) alleles (Δ*osrP* and *osrP::FLAG*, respectively), we implemented a double crossover recombination strategy using the pNPTS138 plasmid (42, 43). Briefly, an in-frame deletion allele of *osrP* was generated using primers listed in **Table S2** in the supplemental material and combined using splice-overlap-extension. The deletion allele carries a 5’ (UP) and 3’ (DOWN) flanking sequences of *osrP* and was ligated in the multiple cloning site (MCS) of a digested pNPTS138 using the restriction enzymes HindIII and SpeI. The tagged allele *osrP::FLAG* was generated using three pieces, two with primers and one with a gene block (Gblock) listed in **Table S2**. The tagged allele was cloned into HindIII and SpeI digested pNPTS138 using Gibson assembly of all three pieces. The first recombination was achieved using a tri-parental mating described in the “Plasmid integration in *C. crescentus*” section with the plasmid-specified antibiotic, kanamycin (5 μg/ml). Single colony exconjugants were inoculated into liquid PYE for 6–16 hours in a rolling 30°C incubator for non-selective growth. Nonselective liquid growth allows for the second recombination event to occur, which either restores the native locus or replaces the native locus with the pNPTS138 insertion sequence. Counter-selection for the second recombination of pNPTS138 was carried out on PYE agar with 3% (w/v) sucrose. This selects for loss of the *sacB* gene during the second recombination event. Colonies were subjected to PCR genotyping and/or sequencing to confirm the allele replacement.

### Genetic complementation constructs in *C. crescentus*

Tandem P1-*gsrN* alleles (overexpression by multiple copies of P1-*gsrN*) were constructed using a Gblock template amplified with three sets of unique primers. Each end of the amplified products contained unique overlap ends for Gibson assembly into pMT552 digested with KpnI and SacI. *gsrN* alleles cloned into the *vanA* locus are antisense to the vanillate inducible *vanA* promoter. An in-frame stop codon was designed at the restriction enzyme/ligation site downstream of the *vanA* promoter to ensure that translational read-through of the *vanA* transcript did not disrupt *gsrN* transcription. Xylose-inducible *osrP* (pMT680-*osrP*) had its entire coding sequence cloned in frame with the start site of *xylX*.

### β-galactosidase reporter constructs

Transcriptional and transcriptional-translational (TT) reporters utilized the replicating plasmids pRKlac290 and pPR9TT, respectively (39, 44). pRKlac290 has a tetracyline resistance marker and pPR9TT has a chloramphenicol resistance marker. Insertion sequences of *osrP* used the primers in **Table S2**. The template for *osrP(RS1)* was created using splice-overlap-extension and the template for *osrP(RS2)* was a gblock. Templates were then amplified with the same primers as the wild-type *osrP* reporters. The transcriptional reporter used the restriction sites EcoRI and HindIII to ligate into pRKlac290. The transcriptional-translational reporter used the restriction sites KpnI and HindIII to ligate into pPR9TT.

### Osmotic stress assay

Liquid cultures were passaged several times before stress treatment to ensure that population growth rate and density were as consistent as possible prior to addition of sucrose (hyperosmotic stress). Briefly, starter cultures were inoculated in liquid M2X medium from colonies grown on PYE-agar plates. Cultures were grown overnight at 30°C in a rolling incubator. Overnight cultures were then diluted back to an optical density reading of OD_660_ = 0.05 and grown in a rolling incubator at 30°C for 7–10 hr. After this period, cultures were re-diluted with M2X to OD_660_ = 0.025 and grown overnight for 16 hr at 30°C in a rolling incubator. After this period, OD_660_ was consistently 0.85–0.90. These cultures were then diluted to OD_660_ = 0.05 and grown for 1 hr and split into two tubes. One tube received sucrose treatment from a liquid stock of 80% (w/v) and the other tube was treated with water. Both cultures were grown for 5 hours in a rolling 30°C post treatment of a final concentration of 300 mM sucrose. This allowed for the dynamic range to compare CFUs from Δ*gsrN*, wild type, and *gsrN*^++^. Treated cultures and untreated cultures were subsequently titered in a 10-fold dilution series (10 μL sample in 90 μL of PYE) in 96-well plates. 5 μL from each dilution were spotted on PYE-agar. Once spots dried, plates were incubated at 30°C for 2 days. Clearly visible colonies begin to form after 36 hours in the incubator.

### Northern Blot

RNA samples were resolved on a urea-denaturing 10% acrylamide: bisacrylamide (29:1), tranferred onto a Zeta-Probe Blotting Membrane with a Trans-Blot^®^ SD Semi-Dry Transfer Cell. Blots were hybridized with a hybridization buffer containing the radiolabeled oligonucleotide probes in a rolling 65°C incubator. Hybridization buffer had a GsrN probe concentration ~1 nM and 5S rRNA probe concentration was ~2 pM. Membranes were then wrapped in plastic wrap and placed directly against a Molecular Dynamics Phosphor Screen. Screens were imaged with Personal Molecular Imager™ (PMI™) System. For detailed buffer recipes and step-by-step instructions refer to (11). Cultures used for the extraction of RNA were passaged in the same manner outlined in the “Osmotic stress assays” section above. Exponential phase cultures were harvested from the last starter (i.e., the OD_660_=0.05 culture at the 16 hour time point) when it reached an OD_660_ of 0.20–0.25. Exponential phase cultures (OD_660_ of 0.20–0.25) harvested for extraction of RNA were pelleted at 15000x g for 3 minutes at ≈23°C (i.e. room temperature) and subjected to a TRIzol extraction (refer to detailed protocol (11)). Radiolabeled oligonucleotides were labeled with T4 PNK (refer to (11) for detailed protocol). Oligonucleotide sequences used for Northern blot probing can be found in **Table S2** in the supplement material.

### RNA-Seq sample preparation and analysis

RNA-Seq samples were extracted using the TRIzol protocol described in (11). For the first RNA-Seq experiment with seven Δ*sigT* (3 stressed and 4 unstressed) and eight WT (4 stressed and 4 unstressed) samples, cells were grown similarly to those described in the “Osmotic stress assay” section. Specifically, liquid M2X cultures were inoculated from PYE agar plates and grown shaking at 200 RPM, 30°C overnight. Cultures were then diluted into fresh M2X to OD_660_ = 0.025 and grown at 200 RPM, 30°C for 18 hours. These overnight cultures were then diluted to OD_660_ = 0.15, and grown for 1 hour at 200 RPM, 30°C before the addition of 150 mM Sucrose (treated) or water (untreated). Samples were grown for 3 hours at 200 RPM, 30°C before TRIzol extractions. Resuspended RNA pellets after the 75% ethanol wash were purified twice by RNeasy Mini Kit column (100 μL sample, 350 μL RLT, 250 μL 100% ethanol). In each iteration, immobilized RNA was subjected to an on-column DNase digestion with TURBO™ DNase for 30 minutes at 30°C with 70 µL DNase Turbo (7 µL DNase, 7µL 10X Buffer, 56µL diH_2_O) before washing and elution. For the second RNA-Seq experiment with 8 *gsrN*^++^ (4 stressed and 4 unstressed) and 6 WT (3 stressed and 3 unstressed) samples, cells were grown as described in the “Osmotic stress assay” section. Specifically, treated cultures were grown for 5 hours in M2X with a final concentration 150 mM sucrose and untreated with water in a rolling 30°C incubator before TRIzol extractions. Resuspended RNA pellets after the 75% ethanol wash were loaded onto an RNeasy Mini Kit column (100 μL sample, 350 μL RLT, 250 μL 100% ethanol). Immobilized RNA was then subjected to an on-column DNase digestion with TURBO™ DNase. DNase treatment was repeated twice on the same column; each incubation was 30 minutes at 30°C with 70 μL solutions of DNase Turbo (7 μL DNase, 7 μL 10x Buffer, 56 μL diH2O). For all RNA-seq samples, after elution from the RNeasy column, rRNA was depleted using Ribo-Zero rRNA Removal (Gramnegative bacteria) Kit (Epicentre). RNA-seq libraries were prepared with an Illumina TruSeq stranded RNA kit according to manufacturer’s instructions. The libraries were sequenced on an Illumina HiSeq 4000 at the University of Chicago Functional Genomics Facility. Analysis of whole genome RNA-seq data was conducted using the CLC Genomics Workbench version 11.0. Reads were mapped to the *C. crescentus* NA1000 genome (accession CP001340.1) (45).

### Soluble protein extraction for LC-MS/MS and analysis

Total soluble protein for proteomic measurements was extracted from cultures passaged similarly to the “Osmotic stress assays” section, except that cultures were subjected to 150 mM sucrose. Cells were spun down at 8000g at 4°C for 15 minutes. Cells were resuspended in 6 mL of ice-cold lysis buffer. Cells were mechanically lysed in LV1 Microfluidizer. Lysate was then spun down at 8000g at 4°C for 15 minutes. Protein samples were resolved on a 12% MOPS buffered 1D Gel (Thermo Scientific) for 10 minutes at 200V constant. Gel was stained with Imperial Protein stain (Thermo Scientific), and a ~2 cm plug was digested with trypsin. Detailed trypsin digestion and peptide extraction by the facility is published in (46). Samples for analysis were run on an electrospray tandem mass spectrometer (Thermo Q-Exactive Orbitrap), using a 70,000 RP survey scan in profile mode, m/z 360–2000 Fa, with lockmasses, followed by 20 MS/MS HCD fragmentation scans at 17,500 resolution on doubly and triply charged precursors. Single charged ions were excluded, and ions selected for MS/MS were placed on an exclusion list for 60s (46). Raw files of LC-MS/MS data were processed using the MaxQuant software suite v1.5.1.2 (47). Samples were run against a FASTA file of proteins from the UniProt database (UP000001364) and standard contaminants. The label free quantitation (LFQ) option was turned on. Fixed modification included carbamidomethyl (C) and variable modifications were acetyl or formyl (N-term) and oxidation (M). Protein group files were created for two comparisons: wild-type (3 samples) versus *gsrN*^++^ (4 samples) untreated and wild-type (3 samples) versus *gsrN*^++^ (4 samples) sucrose-treated. LFQ values for each proteingroup.txt file were extracted for analysis. Average LFQ values were only calculated if 2 or more LFQ values were found for wild-type samples and if 3 or more LFQ values were found for *gsrN*^++^ samples. This allowed for protein groups that had a sufficient amount of signal across all the samples and analyses to be considered for comparison. Once averages for each protein group were calculated, we calculated the fold change between samples from different backgrounds by dividing the averages and taking the log-2 transformation, *log_2_*(Fold). Multiple t-tests were conducted using the LFQ criteria described previously. We used the multiple t-test analysis from GraphPad Prism version 7.0 for MacOS, GraphPad Software, La Jolla California USA, www.graphpad.com. The false discovery rate (Q) value was set to 5.000% and each row was analyzed individually, without assuming a consistent SD.

### σ^T^-binding site search

A binding site search was conducted on negative differentially regulated genes identified in the RNA-Seq study (i.e. genes downregulated in Δ*sigT* relative to wild type; fold change ≤ −1.5 and FDR ≤ 0.05) (**Table S3**). From this set of genes, we organized all genes into operon units based on the DOOR database (48, 49); however, we only put a gene into the context of an operon if the leading gene in the operon was also in the core *sigT* regulon. We then took the lead genes for each operon and searched 250 nucleotides upstream of the annotated coding start site. These windows were then scanned for the degenerate σ^T^-binding site combinations described in **Fig. S2**.

### 5’ rapid amplification of cDNA ends (RACE)

Rapid amplification of cDNA 5’ends of GsrN was carried out using components of the FirstChoice RLM-RACE Kit. Cloning of cDNA library was carried out with the Zero Blunt TOPO PCR Cloning Kit. Total RNA from *gsrN*^++^ strains was extracted from stationary phase cultures (OD660 = 0.95–1.0) as described in the “Northern Blot” section. Briefly, 10 μL Tobacco Acid Pyrophosphatase (TAP) reactions used 5 μg of total RNA with 2 μL of TAP and 1 μL of TAP buffer with remaining volume comprised of Nuclease-free water. Reactions were incubated at 37°C for 1 hour. TAP-treated samples were then subjected to ligation in parallel with no-TAP total RNA samples. Tap RNA sample ligation reactions (10 μL) follow: 2 μL of TAP treated RNA, 1 μL of 5’RACE adaptor, 1 μL of T4 RNA Ligase, 1 μL 10X T4 RNA Ligase Buffer, and 4 μL Nuclease-free water. No-TAP RNA sample ligation reactions (10 μL) follow: 3 μg of untreated total RNA, 1 μL of 5’RACE adaptor, 1 μL of T4 RNA Ligase, 1 μL 10X T4 RNA Ligase Buffer, and remaining volume of Nuclease-free water. Reactions were incubated at 37°C for 1 hr. For the reverse transcription reaction (first strand synthesis), we used the random dodecamer provided in the kit, as well as, the M-MLV Reverse transcriptase and used the recommended reaction volumes in the kit. Reaction was incubated at 42°C for 1 hour. Samples were then kept frozen in a −20°C freezer. For second strand synthesis and amplification, we used KOD Hot Start DNA Polymerase with the 5’RACE inner primer complementary to the adapter and an *osrP*-specific primer 380 nucleotides away from the coding start site **(Table S2)**. The 25 μL reactions follow: 12.5 μL 2X Buffer, 0.5 μL KOD Polymerase, 5 μL of 2 mM dNTP, 2.5 μL of 50% DMSO, 1.5 μL of 5 mM forward primer, 1.5 μL of 5 mM reverse primer, and 1.5 μL of reverse transcribed 1st strand synthesis cDNA. Reaction protocol follows: 3 min 95°C incubation, followed by a 35-cycle reaction consisting of a 15 s 95°C melting step, a 15 s 60°C annealing step, a 30 s 68°C extension step, and a final 1 min 68°C extension step. PCR products were blunt-cloned using the Zero Blunt TOPO PCR Cloning Kit. First, a 5 μL pre-reaction mix consisting of 2 μL PCR product, 1 μL kit salt solution, and 2 μL water was prepared. 1 μL of the pCR-Blunt II-TOPO was then added to the pre-reaction mix and incubated at room temperature for 5 min and then immediately put on ice. Ligation reaction was then incubated with ice-thawed chemically competent *E. coli* cells for 5 min. Cells were heat shocked for 30 s at 42°C, then incubated on ice for 5 min. 250 μL of SOC media was then added to the cells and incubated 37°C in a shaking incubator. Fifty microliters of outgrown cells were placed on LB-Kanamycin plates with an antibiotic concentration of 50 μg/mL. Single colonies were grown overnight and sequenced with an internal *osrP* specific primer that maps 300 nucleotides from the annotated coding start and M13R primers **(Table S2)**. Sequences were submitted to the University of Chicago Comprehensive Cancer Center DNA Sequencing and Genotyping Facility. Chromatograph traces were analyzed with Geneious 11.0.2. Traces were subjected to mapping and trimming of the 5’RACE inner primer/adaptor sequence and the flanking regions used for blunt-cloning.

### β-galactosidase assay

To assess reporter gene expression, liquid cultures were passaged several times as described in the “Osmotic stress assay” section above. However, cultures were placed in a 30°C shaker instead of a 30°C rolling incubator. Exponential phase cultures were taken from the OD_660_ = 0.05 culture made from the 16 hr overnight culture and split when an OD_660_ ~.09–0.1 was reached. One split culture was treated to a final concentration of 150 mM sucrose and the other with the equal volume of water. Stress and unstressed cultures were then grown for three hours in a 30°C shaker and then harvested. β-galactosidase activity from chloroform-permeabilized cells was measured using the colorimetric substrate o-nitrophenyl-b-D-galactopyranoside (ONPG). 1 mL enzymatic reactions contained 350 μL of chloroform-permeabilized cells, 550 μL of Z-buffer (60 mM Na_2_HPO_4_, 40 mM, NaH_2_PO_4_, 10 mM KCl, 1 mM MgSO_4_), and 200 μL of 4 mg/mL ONPG in 0.1 M KPO_4_, pH 7.0. Chloroform-permeabilized cell samples were prepared from 150 μL of culture, 100 μL of PYE, and 100 μL of chloroform (chloroform volume is not included in the final calculation of the 1 mL reaction). Chloroform-treated cells were vortexed for 5–10 seconds to facilitate permeabilization. Z buffer and ONPG were added directly to chloroform-permeabilized cells. Reactions were incubated in the dark at room temperature and quenched with 1 mL of 1 M Na_2_CO_3_. Each reporter construct was optimized with different reaction times empirically determined by the development of the yellow ONPG pigment. Miller units were calculated as:

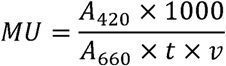

A_420_ is the absorbance of the quenched reaction measured at 420 nm on a Spectronic Genesys 20 spectrophotometer (ThermoFisher Scientific, Waltham, MA). A_660_ is the optical density of the culture of cells used for the assay. *t* is time in minutes between the addition of ONPG and the quenching with Na_2_CO_3_. *v* is the volume in milliliters of the culture added to the reaction.

### Western Blot

Strains from which protein samples were prepared for Western blot analysis were grown and passaged as outlined in the “Osmotic stress assays” section; however, cultures were grown to an OD_660_=0.25-0.30, split, and treated with 150 mM sucrose for 3.5 hours. This change allowed for detection of *osrP::FLAG* signal in untreated *gsrN*^++^ cultures and treated wild-type cultures. 4.5 mL of these cultures was then pelleted, resuspended in 100 μL of Western blot buffer (10 mM Tris pH 7.4, 1 mM CaCl_2_, and 5 μg/mL of DNase), and mixed with 100 μL SDS-Loading buffer. Samples were boiled at 85°C for 10 minutes, and 25–30 μL of each sample was loaded onto a Mini-PROTEAN TGX Precast Gradient Gel (4–20%) with Precision Plus Protein™ Kaleidoscope™ Prestained Protein Standards. Samples were resolved at 35 mA constant current in SDS running buffer (0.3% Tris, 18.8% Glycine, 0.1% SDS). Gels were run until the 25 kDa marker reached the bottom of the gel. Gel was transferred to an Immobilon^®^-P PVDF Membrane using a Mini Trans-Blot^®^ Cell after preincubation in Western transfer buffer (0.3% Tris, 18.8% Glycine, 20% methanol). Transfer was carried out at 4°C, 100 V for 1 hour and 20 minutes in Western transfer buffer. The membrane cut into two pieces right above the 50kD marker. Top half was stained with Coommassie Brilliant Blue for 10 minutes, washed with 45% Ethanol and 10 % Acetic acid, and then washed again with 90% Ethanol 10% Acetic acid. Upon destaining, image was taken with a ChemiDoc MP Imaging System version 6.0. Bottom half was blocked in 5% (w/v) powdered milk in Trisbuffered saline with tween (TBST: 137 mM NaCl, 2.3 mM KCl, 20 mM Tris pH 7.4, 0.1% (v/v) Tween 20) overnight at room temperature on a rotating platform. Primary incubation with an anti-DYKDDDDK Monoclonal Antibody (clone FG4R) was carried out for 3 hours in 5% powdered milk TBST at room temperature on a rotating platform (4 μL antibody in 12 mL). Membrane was then washed 3 times in TBST for 5 minutes each at room temperature on a rotating platform. Secondary incubation with Goat anti-Mouse IgG (H+L) Secondary Antibody, HRP was for 1 hour at room temperature on a rotating platform (3 μL antibody in 15 mL). Finally, membrane was washed 3 times in TBST for 10 minutes each at room temperature on a rotating platform. Chemiluminescence was performed using the SuperSignal™ West Femto Maximum Sensitivity Substrate and was imaged using a ChemiDoc MP Imaging System version 6.0. Chemiluminescence was measured using the ChemSens program with an exposure time of ~2.5 minutes.

### Accession number(s)

RNA-Seq data are available in the NCBI GEO Database under accession GSE114971. LC-MS/MS data is available in the PRIDE proteomic database under accession PXD010072.

## Acknowledgements

We thank Aretha Fiebig for constructive feedback on this manuscript during the revision process. This work was funded by NIH award R01GM087353 to S.C. M.Z.T. was supported by an NSF Graduate Research Fellowship, and B.J.S. is supported by NIH award F32GM128283.

